# Time distributed data analysis by Cosinor.Online application

**DOI:** 10.1101/805960

**Authors:** Lubos Molcan

## Abstract

Disturbed biological oscillations often represent pathology and thus have a prognostic character. The most studied are 24-h (circadian) and shorter (ultradian) oscillations from them. A cosinor analysis often evaluates the presence and significance of circadian and ultradian rhythms. Skilled researchers can use MATLAB, R, Python, or other programming languages, while those less experienced often use outdated applications that require a specific operating system version or outdated add-ons. Therefore, we developed CosinorOnline, a simple web-based application coded in PHP and JavaScript to evaluate the presence and significance of different oscillations. Users can set the period length on the application’s page and insert their data. The results consist of a numerical evaluation and an adjustable plot. There is also a unique identifier to reload or immediately delete data analysis within one month. After this period, all data are automatically deleted from the app. We compared the functionality of CosinorOnline with Cosinor2 (R package) and Chronos-Fit (Windows app). The 24-h variability analysis was identical for all three applications. The evaluation of ultradian variability was the same for CosinorOnline and Cosinor2 and slightly different for Chronos-Fit. CosinorOnline and Chronos-Fit result in acrophase in units of time (decimal form), while Cosinor2 is in radians. In conclusion, CosinorOnline is a simple, easy-to-use web application to inspect time data that provides reliable results without additional installation and runs in modern web browsers. The application does not track users and aims to help users quickly decide whether their data is suitable for more profound analysis using cosinor analysis.

## 1 Introduction

In living systems, processes vary over time because of the integration of biological and environmental interactions. According to the period length, we divide rhythms into ultradian (period < 20 hours), circadian (period 20 – 28 hours) and infradian (period > 28 hours) domains [1,2]. The most studied are circadian rhythms, prominent in physiological systems during normal light/dark conditions [3]. In a short-term manner, shifts work, dimmed artificial light during the night, jet lag, and stress significantly affect the course and presence of circadian variability of physiological parameters [3–9]. Moreover, the long-term disturbance of the circadian oscillations can change system-level set points and lead to pathological processes [10–12], including cancer development [13].

The presence and significance of circadian oscillations in rhythmic data can be estimated using cosinor analysis [14]. Cosinor analysis not only determines whether the data oscillate significantly but also describes the rhythmic data using: 1) mesor, the average value around which the variable oscillates; 2) amplitude, the difference between the peak and the mean value; and 3) acrophase, the time of peak activity, from a simple cosine function [15].

The simple cosine function can be evaluated with a wide range of locally installed native applications that do not always run intuitively and often require a specific operating system version or the installation of outdated add-ons. Skilled researchers can use the MATLAB, R or Python programming languages to adjust the cosinor code, change the input form of the time series (radians, degrees, or hours) or modify estimated period lengths and other parts of the cosinor model [16–19]. Less skilled researchers often use unsupported outdated software, whereas each works with different input file types and formats. On the other hand, server-based software applications are popular in biomedicine [20–22] because they are straightforward and run in standard web browsers 24/7. Therefore, we developed a simple web-based application (CosinorOnline) to evaluate the presence and significance of oscillations in rhythmic data using only a web browser.

## 2 Material and methods

### 2.1 Mathematical modelling and code writing

We programmed the application in PHP and JavaScript in line with the MATLAB code [18], which implements a standard cosinor method [15]. In the beginning, we solved a system of linear equations (Equation 1). Because of PHP implementation, we used matrix inversion instead of Gauss-Jordan elimination. This approach can provide false results if the matrix is poorly conditioned in extreme cases.

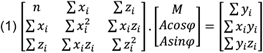

From Equation 1, where *x_i_* = *cos*(*ωt_i_*), *z_i_* = *sin*(*ωt_i_*), *y_i_* is the value of series at time *t*, and *n* is a count of input values, we obtained mesor (*M*), *Acosφ* and *Asinφ* from which we calculated amplitude (*A*) (Equation 2). Using inverse tangent in radians and the signum function, we estimated acrophase (*φ*) and converted *φ* from radians to degrees and hours (Equation 3)

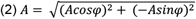

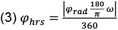

Mesor, amplitude and acrophase were entered into Equation 4, where *ω* is the period length specified by a user, and we did cosine fit to the imported data.

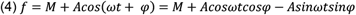

The significance of model fitting was done using the p-value as the F probability density function (F-value).

Input and output data are handled and temporarily stored in a MySQL database. Storing data in the database is only a technical implementation to allow users to reload previous analyses on different devices at various times over one month. Any other data (e.g., operating system, browser, country, and other) and cookies are not tracked and stored by our application or other third-party tracking tools.

The MySQL database loads data using a randomly generated unique identifier string. Original input data are visualized as a scatter plot with a cosine fit as a line plot. Due to online usage, cross-platform and cross-browser compatibility, we used Google Charts [23], which are easy to use for online visualization of a wide range of data sets.

### 2.2 Software usage

Software codes are hosted on the encrypted domain, and the application is visualized by HTML and CSS (Bootstrap) at the https://cosinor.online address. Users can enter the period length, time, and data on the main page in decimal form. Thus, 8:00 a.m. is 8.00, 8:15 a.m. is 8.25, 8:00 p.m. is 20.00, 8:15 p.m. is 20.25 and so forth (**Figure 1**). Time and data columns must be the same size.

**Figure 1:**
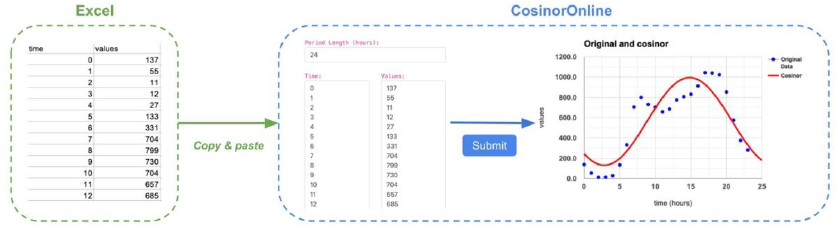
Loading of data to the CosinorOnline app. Users can copy and paste data from an Excel file to the web form in decimals.

After submitting the data, users obtain outputs from the model describing the selected period length (**Figure 2**). The application calculates mesor and amplitude in the units of the user’s data; acrophase is provided in hours, decimal form because hours compared to radians are more understandable for ordinary users. The chart’s title and vertical and horizontal axes names and intervals are adjustable. Charts and output data are downloadable.

**Figure 2:**
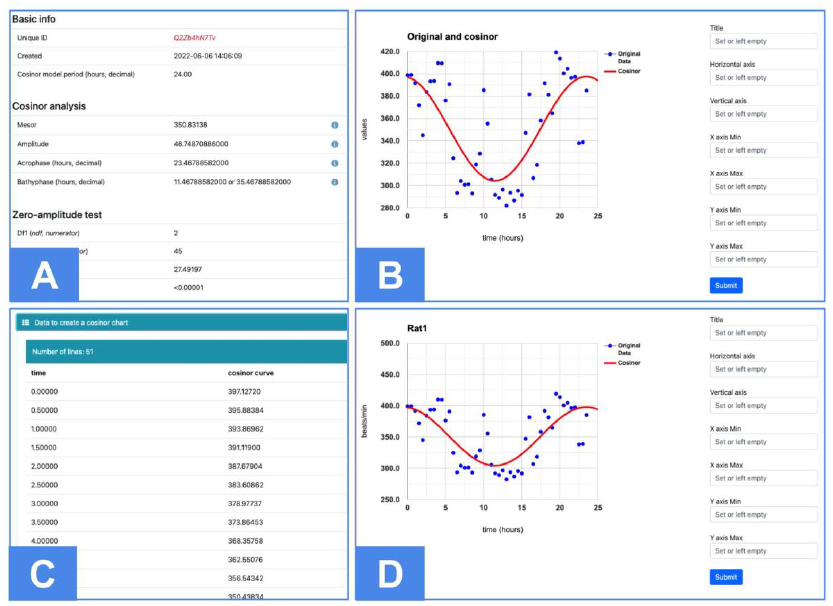
(A) On the output page, users can find information about the date and time of data analysis and the unique identifier string to reopen or remove results. Mesor and amplitude are in the units of the user’s input data. Acrophase and bathyphase are provided in hours, decimal form. F- and p-value determines whether the selected model period significantly fits user data. (B) Original and cosinor data are plotted on the chart. (C) Users can create charts themselves from the calculated data of the cosinor model (D) or edit the chart directly on the web page.

Users can also find information about the date and time of the analysis and the unique identifier string on the output page. A unique identifier string allows users to reopen or remove their data and results over one month.

### 2.3 Comparison with other applications

Many applications evaluate cosinor analysis, but many require installation on a local drive, a specific type/version of the operating system, or the presence of various addons. Therefore, we compared our CosinorOnline application only with Cosinor2 (R package; [17]) and Chronos-Fit [24], a Windows app, to which professor B. Lemmer gently provided us with a license key to activate it.

As input data (Supplementary Table 1), we analyzed the heart rate of telemetrically measured rats (R1: 325 g; R2: 286 g; R3: 342 g; R4: 285 g) and compared outputs (mesor, amplitude, acrophase and the F-value) from CosinorOnline, Cosinor2 and Chronos-Fit applications in two ways. 1) We evaluated the presence and significance of the 24-h period in data sets with 24 (Data set 1), 48 (Data set 2) and 96 (Data set 3) values. 2) In Data set 3 (96 values), we analyzed the presence of the 24-h and ultradian (16-, 14-, 13-, 9-, 8- and 3-h) periods.

### 2.4 Data analysis

Data were statistically compared by one-way ANOVA and visualized in the statistical software R, version 3.6.3 [25] or the CosinorOnline application.

## 3 Results

### 3.1 Differently sized data sets

Firstly, we estimated 24-h periods in data sets of different sizes (**Figure 3**). Cosinor2 incorrectly calculated acrophase by the basic Rhythm Detection Test function (cosinor.detect(x)); thus, we had to calculate the acrophase separately using the correct.acrophase(x) function in Cosinor2. In contrast, Chronos-Fit converted acrophase to hours (in decimal form) natively, like our web-based CosinorOnline application. Chronos-Fit, CosinorOnline and Cosinor2 provided identical results in detecting 24-h periods (**Table 1**; Supplementary Table 2).

**Figure 3:**
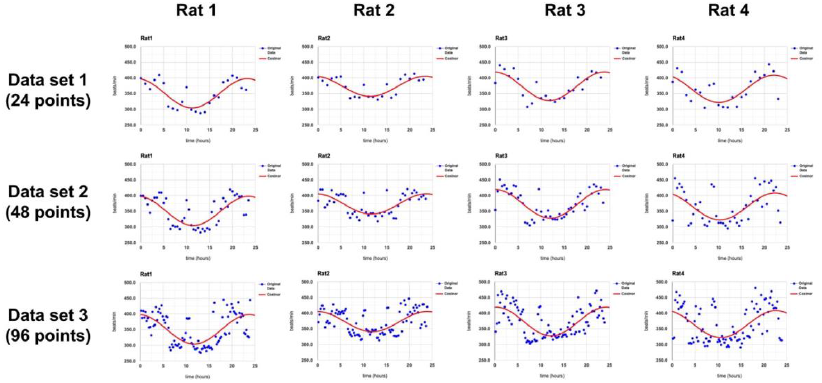
The CosinorOnline application fits a 24-h period (red line) into three differently sized data sets of individual rats’ heart rates. The original data are visualized as blue dots.

**Table 1:**
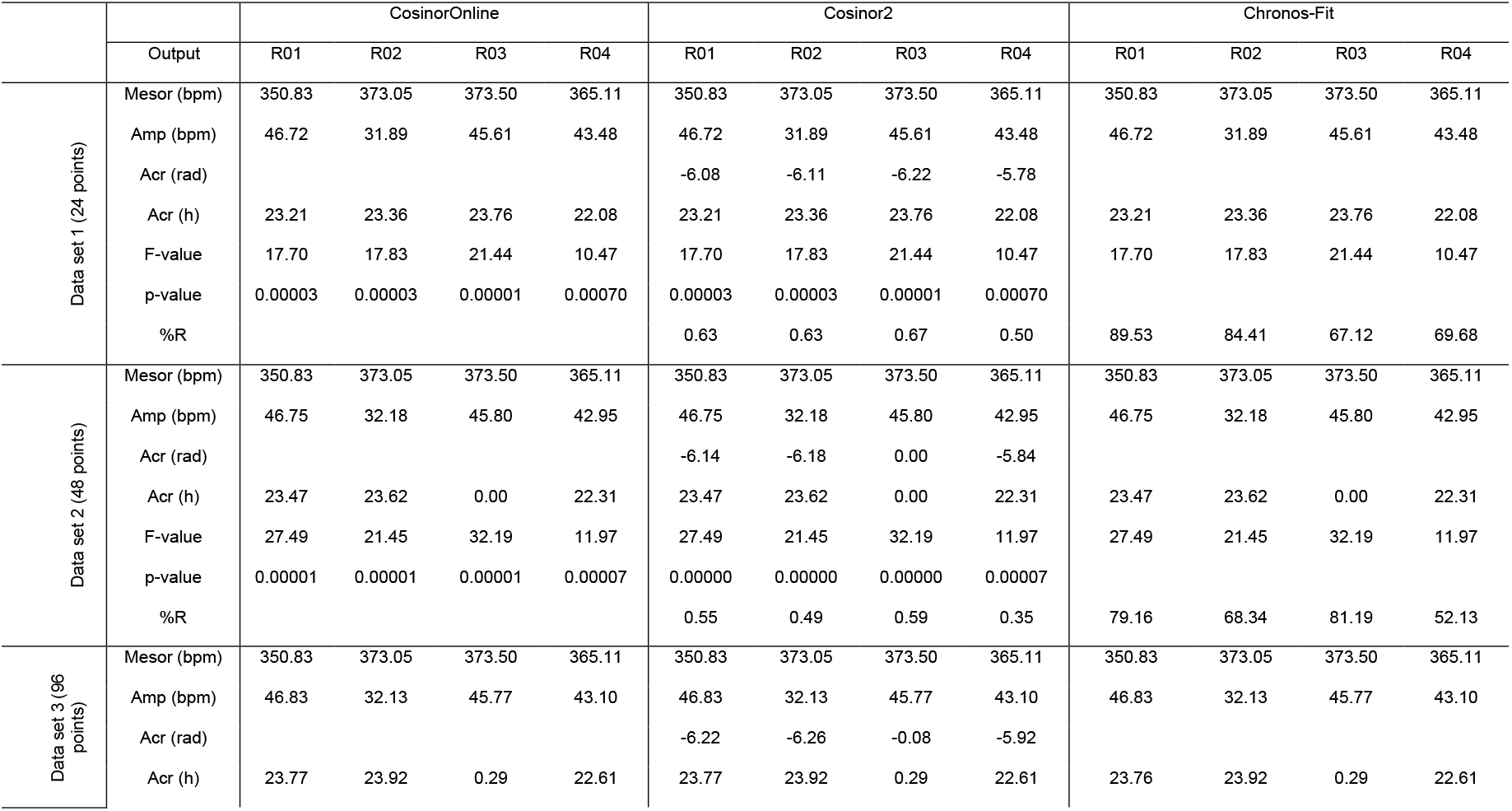
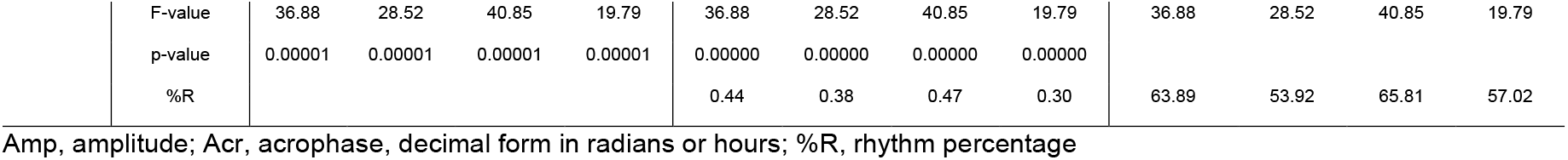
Comparison of 24-h periods of individual rat’s heart rate analysed by CosinorOnline, Cosinor2 and Chronos-Fit.

|  |  | CosinorOnline |  |  |  | Cosinor2 |  |  |  | Chronos-Fit |  |  |  |
| --- | --- | --- | --- | --- | --- | --- | --- | --- | --- | --- | --- | --- | --- |
|  | Output | R01 | R02 | R03 | R04 | R01 | R02 | R03 | R04 | R01 | R02 | R03 | R04 |
| Data set 1 (24 points) | Mesor (bpm) | 350.83 | 373.05 | 373.50 | 365.11 | 350.83 | 373.05 | 373.50 | 365.11 | 350.83 | 373.05 | 373.50 | 365.11 |
|  | Amp (bpm) | 46.72 | 31.89 | 45.61 | 43.48 | 46.72 | 31.89 | 45.61 | 43.48 | 46.72 | 31.89 | 45.61 | 43.48 |
|  | Acr (rad) |  |  |  |  | -6.08 | -6.11 | -6.22 | -5.78 |  |  |  |  |
|  | Acr (h) | 23.21 | 23.36 | 23.76 | 22.08 | 23.21 | 23.36 | 23.76 | 22.08 | 23.21 | 23.36 | 23.76 | 22.08 |
|  | F-value | 17.70 | 17.83 | 21.44 | 10.47 | 17.70 | 17.83 | 21.44 | 10.47 | 17.70 | 17.83 | 21.44 | 10.47 |
|  | p-value | 0.00003 | 0.00003 | 0.00001 | 0.00070 | 0.00003 | 0.00003 | 0.00001 | 0.00070 |  |  |  |  |
|  | %R |  |  |  |  | 0.63 | 0.63 | 0.67 | 0.50 | 89.53 | 84.41 | 67.12 | 69.68 |
| Data set 2 (48 points) | Mesor (bpm) | 350.83 | 373.05 | 373.50 | 365.11 | 350.83 | 373.05 | 373.50 | 365.11 | 350.83 | 373.05 | 373.50 | 365.11 |
|  | Amp (bpm) | 46.75 | 32.18 | 45.80 | 42.95 | 46.75 | 32.18 | 45.80 | 42.95 | 46.75 | 32.18 | 45.80 | 42.95 |
|  | Acr (rad) |  |  |  |  | -6.14 | -6.18 | 0.00 | -5.84 |  |  |  |  |
|  | Acr (h) | 23.47 | 23.62 | 0.00 | 22.31 | 23.47 | 23.62 | 0.00 | 22.31 | 23.47 | 23.62 | 0.00 | 22.31 |
|  | F-value | 27.49 | 21.45 | 32.19 | 11.97 | 27.49 | 21.45 | 32.19 | 11.97 | 27.49 | 21.45 | 32.19 | 11.97 |
|  | p-value | 0.00001 | 0.00001 | 0.00001 | 0.00007 | 0.00000 | 0.00000 | 0.00000 | 0.00007 |  |  |  |  |
|  | %R |  |  |  |  | 0.55 | 0.49 | 0.59 | 0.35 | 79.16 | 68.34 | 81.19 | 52.13 |
| Data set 3 (96 points) | Mesor (bpm) | 350.83 | 373.05 | 373.50 | 365.11 | 350.83 | 373.05 | 373.50 | 365.11 | 350.83 | 373.05 | 373.50 | 365.11 |
|  | Amp (bpm) | 46.83 | 32.13 | 45.77 | 43.10 | 46.83 | 32.13 | 45.77 | 43.10 | 46.83 | 32.13 | 45.77 | 43.10 |
|  | Acr (rad) |  |  |  |  | -6.22 | -6.26 | -0.08 | -5.92 |  |  |  |  |
|  | Acr (h) | 23.77 | 23.92 | 0.29 | 22.61 | 23.77 | 23.92 | 0.29 | 22.61 | 23.76 | 23.92 | 0.29 | 22.61 |

|  |  |  |  |  |  |  |  |  |  |  |  |  |
| --- | --- | --- | --- | --- | --- | --- | --- | --- | --- | --- | --- | --- |
| F-value | 36.88 | 28.52 | 40.85 | 19.79 | 36.88 | 28.52 | 40.85 | 19.79 | 36.88 | 28.52 | 40.85 | 19.79 |
| p-value | 0.00001 | 0.00001 | 0.00001 | 0.00001 | 0.00000 | 0.00000 | 0.00000 | 0.00000 |  |  |  |  |
| %R |  |  |  |  | 0.44 | 0.38 | 0.47 | 0.30 | 63.89 | 53.92 | 65.81 | 57.02 |

Amp, amplitude; Acr, acrophase, decimal form in radians or hours; %R, rhythm percentage

### 3.2 Ultradian variability detection

Chronos-Fit, CosinorOnline and Cosinor2 calculated identical mesor, amplitude, acrophase and F-value when detecting 24-h periods (**Table 2**; **Table 3**; Supplementary Table 2; **Figure 4**a). On the other hand, when we tested the presence of ultradian (16-, 14-, 13-, 9-, 8- and 3-h) periods, results from Chronos-Fit slightly differ from CosinorOnline and Cosinor2 (**Table 2**; **Table 3**; Supplementary Table 2; **Figure 4**a).

**Figure 4:**
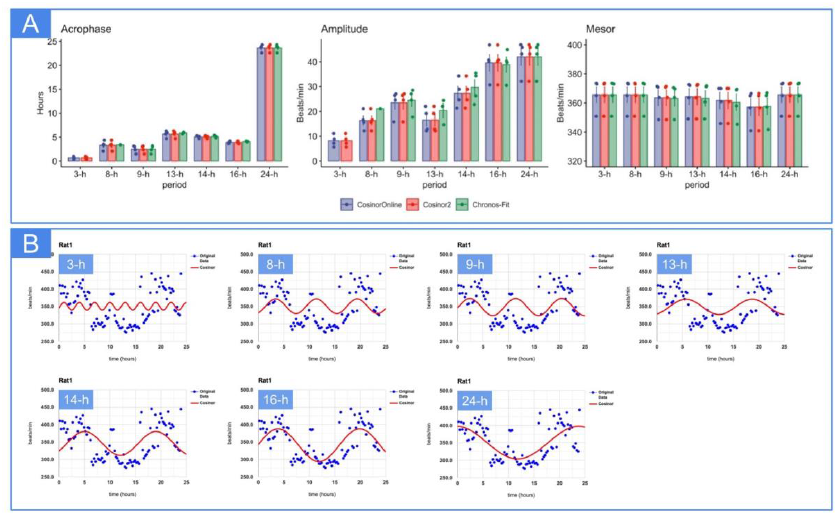
(A) Comparison of acrophase, amplitude and mesor of calculated Data Set 3 (96 points) in CosinorOnline, Cosinor2 and Chronos-Fit. Data are represented as the arithmetic mean and standard error of the mean. (B) The CosinorOnline app fits ultradian periods into Data set 3 of Rat 1.

**Table 2:** Averages of mesor, amplitude (Amp), acrophase (Acr) and F-value from Data set 3 (96 values per each rat) analysed by CosinorOnline, Cosinor2 and Chronos-Fit. Chronos-Fit did not provide (NA) amplitude, acrophase and F-value if F-value was less.

|  |  | 24-h | 16-h | 14-h | 13-h | 9-h | 8-h | 3-h |
| --- | --- | --- | --- | --- | --- | --- | --- | --- |
| CosinorOnline | Mesor (bpm) | 365.62 | 357.25 | 361.68 | 364.35 | 363.63 | 365.62 | 365.62 |
|  | Amp (bpm) | 41.96 | 39.60 | 27.28 | 16.57 | 23.51 | 16.39 | 8.22 |
|  | Acr (h) | 23.65 | 3.90 | 5.04 | 5.66 | 2.50 | 3.31 | 0.68 |
|  | F-value | 31.51 | 22.59 | 8.07 | 2.89 | 6.19 | 3.03 | 0.83 |
| Cosinor2 | Mesor (bpm) | 365.62 | 357.25 | 361.68 | 364.35 | 363.63 | 365.62 | 365.62 |
|  | Amp (bpm) | 41.96 | 39.60 | 27.28 | 16.57 | 23.51 | 16.39 | 8.22 |
|  | Acr (h) | 23.65 | 3.90 | 5.04 | 5.66 | 2.50 | 3.31 | 0.68 |
|  | F-value | 31.51 | 22.59 | 8.07 | 2.89 | 6.19 | 3.03 | 0.83 |
| Chronos-Fit | Mesor (bpm) | 365.62 | 357.74 | 360.42 | 363.17 | 363.13 | 365.62 | 365.62 |
|  | Amp (bpm) | 41.96 | 38.85 | 29.70 | 20.36 | 24.50 | 21.08 | NA |
|  | Acr (h) | 23.65 | 4.08 | 5.11 | 5.90 | 2.51 | 3.39 | NA |
|  | F-value | 31.51 | 22.59 | 8.07 | 4.54 | 6.61 | 4.58 | NA |

**Table 3:** Standard deviations of mesor, amplitude (Amp), acrophase (Acr) and F-value from Data set 3 (96 values per each rat) analysed by CosinorOnline, Cosinor2 and Chronos-Fit. Chronos-Fit did not provide (NA) amplitude, acrophase and F-value if F-value.

|  |  | 24-h | 16-h | 14-h | 13-h | 9-h | 8-h | 3-h |
| --- | --- | --- | --- | --- | --- | --- | --- | --- |
| CosinorOnline | Mesor (bpm) | 10.59 | 11.74 | 11.37 | 10.97 | 10.63 | 10.59 | 10.59 |
|  | Amp (bpm) | 6.74 | 6.67 | 5.57 | 4.83 | 5.24 | 3.66 | 2.34 |
|  | Acr (h) | 0.72 | 0.23 | 0.26 | 0.69 | 0.75 | 0.94 | 0.29 |
|  | F-value | 9.35 | 6.70 | 2.72 | 1.27 | 1.46 | 1.06 | 0.54 |
| Cosinor2 | Mesor (bpm) | 10.59 | 11.74 | 11.37 | 10.97 | 10.63 | 10.59 | 10.59 |
|  | Amp (bpm) | 6.74 | 6.67 | 5.57 | 4.83 | 5.24 | 3.66 | 2.34 |
|  | Acr (h) | 0.72 | 0.23 | 0.26 | 0.69 | 0.75 | 0.94 | 0.29 |
|  | F-value | 9.35 | 6.70 | 2.72 | 1.27 | 1.46 | 1.06 | 0.54 |
| Chronos-Fit | Mesor (bpm) | 10.59 | 11.46 | 11.37 | 10.67 | 10.16 | 10.59 | 10.59 |
|  | Amp (bpm) | 6.74 | 6.03 | 6.16 | 5.12 | 4.79 | NA | NA |
|  | Acr (h) | 0.73 | 0.10 | 0.23 | 0.20 | 0.77 | NA | NA |
|  | F-value | 9.35 | 6.69 | 2.72 | 0.75 | 0.88 | NA | NA |

## 4 Discussion

Biological data vary in time with different period lengths. While some oscillations result from random processes or noise, many fluctuations reflect internal control in living systems [26]. Disrupted or lost oscillations are often accompanied by pathological changes and therefore have a prognostic character. Evaluation of oscillations in biological data is, therefore, essential. Cosinor analysis assesses whether the measured data show significant variability within a defined period length in data sets with a high signal-to-noise ratio [14].

Evaluation of rhythmic data can be done using various locally installed [19] and online [27] applications, allowing analyzing the data in the modern web browsers of different devices, such as computers, tablets or phones. Our web-based CosinorOnline application provides cross-platform and cross-browser compatibility and data availability 24/7 due to PHP, JavaScript and MySQL [28,29]. PHP and MySQL reduce the performance requirements of the user’s device because the calculations are entirely done on the server-side [30]. In addition, MySQL allows researchers to reload the analysis results securely and repeatedly using an original, randomly generated unique identifier string. Some advanced, sophisticated and online apps are on the market [27], but they do not allow data storage (for free) compared to CosinorOnline. Compared to natively installed applications, online applications must also ensure user safety and the potential misuse of uploaded data and user information. Therefore, our application does not use cookies to function, does not use any tracking tools and does not collect information about users in any other way. Only uploaded data and analysis results are temporarily stored in the MySQL database as a free service for users to retrieve results. However, data in the database older than one month are automatically and permanently removed without backups and recovery options.

We simplified the application usage as much as possible. For example, data importing is very straightforward; 1) copy and paste and 2) the acrophase is in hours (decimal form) and not radians or degrees compared to Cosinor2 [17] and some web applications [27], which require files with specific extensions and data formatting. Moreover, in our application, the terms in the result include a brief description. On the output page, users can also find their input data (scatter plot) and the calculated cosine curve (line plot) to better understand the parameters describing their rhythmic data’s behaviour. The titles and intervals of the plot are adjustable, even when reopened. Plots, as well as output data from the cosinor model, can be downloaded by users.

The results from CosinorOnline are identical to those from Cosinor2, even though we changed the period length to ultradian. In contrast, Chronos-Fit gave similar results only for the 24-h period, probably because Chronos-Fit is focused on evaluating mainly the circadian (approximately 24-h) rhythms. However, since we have not seen the application’s source code, it is only our assumption. Cosinor2, in comparison to CosinorOnline, allows, in addition to the fundamental rhythmometry analysis, the comparison of cosinor parameters of two populations while comparing the mesors, amplitudes and acrophases of the two data sets. Like Chronos-Fit, Cosinor2 allows the calculation of rhythm percentage, which calculates the relative force of the selected oscillation [24]. However, the rhythm percentage differs when comparing Cosinor2 and Chronos-Fit (**Table 1**, Supplementary Table 2). Chronos-Fit does not give amplitude and acrophase if the F-value is above the significance limit, which can be adjusted in the application. Chronos-Fit is a relatively robust and stable application. However, its limitation is that the authors had not provided a license key to activate the app for several years. In addition, Chronos-Fit is written for Windows and does not allow crossplatform work compared to multiplatform Cosinor2, which is written in R that is available for Windows, Mac and Linux, and CosinorOnline, which needs only a web browser.

CosinorOnline has been available since 2018 and is actively used by professionals, as evidenced by many journal references [31–34]. The application has performed almost 50,000 analyses according to the number of autoincrement values. Due to the web application does not use any cookies or user tracking tools, we do not know how users used the application sort by year or region.

Limitations. We compared CosinorOnline only with two applications. However, both are commonly used [35,36]. Due to maximizing app efficiency, the cosinor curve in charts could not be completely smooth. Due to PHP implementation, our application can provide false results in extreme cases. CosinorOnline offers only a basic cosinor analysis and does not provide confidence intervals of the mesor, acrophase and amplitude, the zero-amplitude test, the comparative rhythmicity analyses, the multicomponent cosinor analysis, and the goodness-of-fit statistics. On the other hand, CosinorOnline allows quick, intuitive and straightforward data importing and analysis to help users decide whether to do deeper cosinor evaluation using another cosinor-based software [17,19,27] or to analyze the data differently.

## Conclusion

The web-based application CosinorOnline (https://cosinor.online) analyses the presence of various periods as good as Cosinor2 and Chronos-Fit but runs in modern web browsers. Compared to Cosinor2, rhythm acrophase is displayed in hours, not radians. The results from CosinorOnline are freely available for one month. After this period, users’ data are automatically and permanently removed, but users can delete their data immediately after analysis. The application does not use any cookies or web tracking and analysis tools. CosinorOnline is a simple web application that aims to help users decide whether their data is suitable for more profound analysis using cosinor analysis.

## Acknowledgement

The author thanks Dr Richard Kollár and Dr Katarina Bod’ová for their mathematical advice and Dr Monika Okuliarová for the idea of developing an easy-to-use application primarily for students.

